# Long-Term Metabolomics Reference Material

**DOI:** 10.1101/2021.03.24.436834

**Authors:** Goncalo J. Gouveia, Amanda O. Shaver, Brianna M. Garcia, Alison M. Morse, Erik C. Andersen, Arthur S. Edison, Lauren M. McIntyre

## Abstract

The use of quality control samples in metabolomics ensures data quality, reproducibility and comparability between studies, analytical platforms and laboratories. Long-term, stable and sustainable reference materials (RMs) are a critical component of the QA/QC system, however, the limited selection of currently available matrix matched RMs reduce their applicability for widespread use. To produce a RM in any context, for any matrix that is robust to changes over the course of time we developed IBAT (**I**terative **B**atch **A**veraging me**T**hod). To illustrate this method, we generated 11 independently grown *E. coli* batches and made a RM over the course of 10 IBAT iterations. We measured the variance of these materials by NMR and showed that IBAT produces a stable and sustainable RM over time. This *E. coli* RM was then used as food source to produce a *C. elegans* RM for a metabolomics experiment. The metabolite extraction of this material alongside 41 independently grown individual *C. elegans* samples of the same genotype, allowed to estimate the proportion of sample variation in pre-analytical steps. From the NMR data, we found that 40% of the metabolite variance is due to the metabolite extraction process and analysis and 60% is due to sample-to-sample variance. The availability of RMs in untargeted metabolomics is one of the predominant needs of the metabolomics community that reach beyond quality control practices. IBAT addresses this need by facilitating the production of biologically relevant RMs and increasing their widespread use.

---

Biological reference materials are needed to compare metabolomics data across multiple instruments, studies and batches. Whenever there are more samples collected than can be processed in a single ‘run’ there is added unwanted variation that, if captured, can be modeled and removed, leading to more powerful tests.^1^ Readily available long-term biologically relevant reference materials (RMs) represent a critical component to achieve reproducibility.^2, 3^ Commercially available RMs and standard reference materials (SRMs) address some of these needs, but can be expensive to purchase, offer limited quantities, matrix diversity, and have an expiration date.^3^ The National Institute of Standards and Technology (NIST) has a long history of producing biofluid-based SRMs to facilitate standardization and improve comparability and reproducibility of analytical measurements. These SRMs are trademarked Certified Reference Materials (CRMs) and specifically designed to provide certified metabolite levels that serve strict objectives (i.e., calibration, method validation, measurement accuracy).^4–6^ Pooled quality control (QC) samples produced from experimental samples are valuable as they capture instrument variation within the experiment, but have limited value in comparing across experiments, or in synthesizing results from large experiments.^7, 8^ The individual variation intrinsic in subjecting biological material to extraction and quantification is not captured by pooled samples or by chemical standards made after extraction. There is a recognized need for matrix-specific stable RMs that can be used to compare data across long-term studies with multiple batches or across different laboratories and instrumention.^9^

Homogeneous and stable materials that are fit for purpose are reference materials (as per the International Vocabulary of Metrology-VIM).^10^ RM does not require a metrologically valid metabolite quantification (certification) and should be straightforward to produce and maintain. For untargeted metabolomics, additional criteria for a RM are important. Namely, it should (i) be made from the same biological matrix as the experimental samples, (ii) have a profile that is as complex as the experimental samples, (iii) be sustainably produced over time and (iv) facilitate the annotation of known and unknown compounds.

The proteomics community devoted substantial effort to the development and application of RMs, which greatly improved standardization and reproducibility in the field.^4, 9^ The metabolomics community has highlighted the need for RMs as part of the development of resources and practices to measure, detect and prevent unwanted pre-analytical and instrumental variation.^2, 3, 5, 8, 11^.

Here we introduce IBAT (**I**terative **B**atch **A**veraging me**T**hod) that can be used to create a stable RM produced over time in any context. The concept is straightforward: multiple small batches of starting material are produced and aliquoted, and then pooled to generate the RM. A stable and long-lasting RM can be generated by repeating the process over time, as illustrated in Figure 1. IBAT results in a RM that (i) is robust to changes over time, (ii) minimizes variance between batches of RM, (iii) can be used over the course of large-scale experiments, (iv) can be made with a small amount of constant effort and smaller storage space, (v) can be applied to any organism or biological matrix of interest and (vi) can be used for evaluation of multiple sources of variation at multiple points in a metabolomics experiment. To illustrate IBAT, we made and characterized a *Caenorhabditis elegans* reference material. *C. elegans* eats bacteria, which is also subject to variation over time, so to make a stable *C. elegans* RM, we first needed to make an *Escherichia coli* RM that can be fed to *C. elegans*. This two-step IBAT shows the flexibility of the approach, and in the Discussion section we outline strategies to apply IBAT to create other RMs of interest to metabolomics researchers.

**Figure 1:**
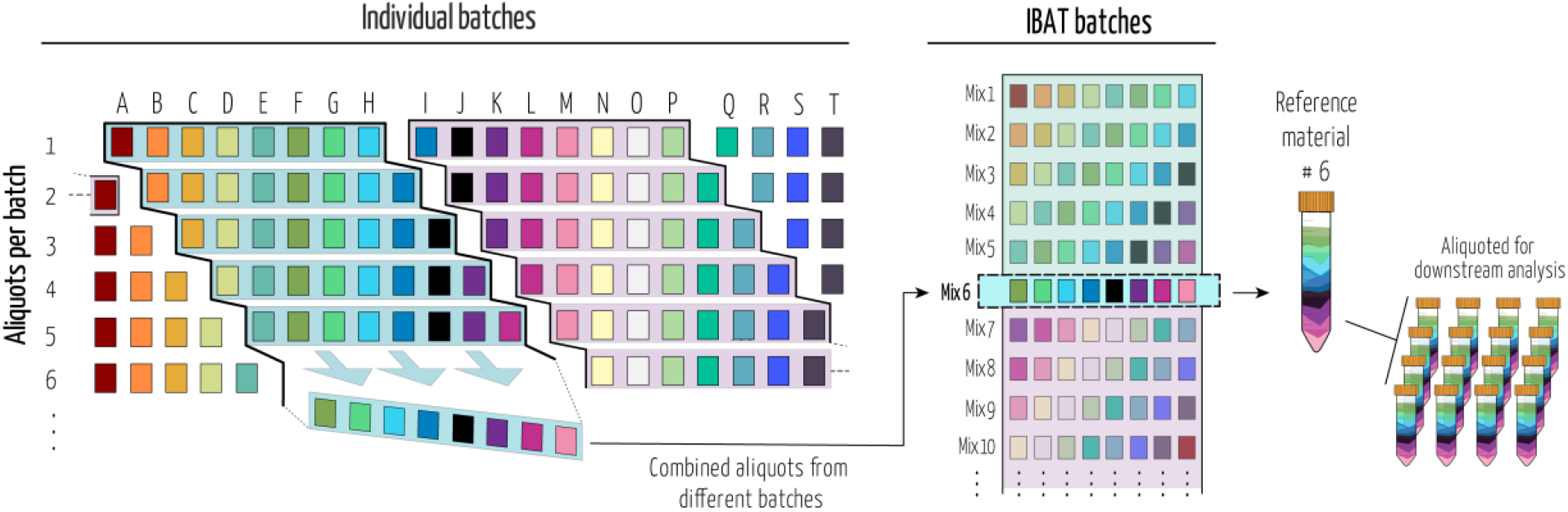
Iterative batch average method (IBAT). Batches of material are represented by columns (same-colored squares and letters). Rows represent homogeneous aliquots from each batch. Examples of sequential batch combinations are rows shaded from blue to purple. Right panel illustrates the IBAT generated pools from individual batches. IBAT is only limited by the number of individual batches produced and can be adjusted to the number of aliquots required and to any material.

## Results

### Production and analysis of an IBAT *E. coli* as a food source for *C. elegans*

For this RM, we used a bioreactor to generate large quantities of bacteria in each batch, but the principle holds on a smaller scale with flasks and a shaker/incubator. We grew 11 different 2 L bioreactor batches (columns in Fig. 1) that each produced an average of 84 g of bacterial paste. Each batch was then aliquoted into 60-90 tubes (rows in Fig. 1) containing 1 g each, with mixing to maintain homogeneity of the aliquots.

We combined single aliquots from five different batches for this *E. coli* RM, such that each tube of IBAT RM contained the same amount of material. The first IBAT sample was made by combining batches A-E (columns in Fig. 1), the second IBAT sample combined batches B-F, etc. When we reached the end of the 11 batches (G-K with 11 batches), the next IBAT sample was made from H-K and an aliquot of A (Fig. 1). A similar IBAT process was applied to *C. elegans*, as described below. We compared the 10 different *E. coli* IBAT samples (Table 1) with individual replicates from all 11 batches.

**Table 1:** List of individual batches pooled together for each stable food source iteration. This process follows the same methodology described in figure 1.

| IBAT *E. coli* iterations | 1 | 2 | 3 | 4 | 5 | 6 | 7 | 8 | 9 | 10 |
| --- | --- | --- | --- | --- | --- | --- | --- | --- | --- | --- |
| Combined individual *E. coli* batches | A to E | B to F | C to G | D to H | E to I | F to J | G to K | H to A | I to B | J to C |

The 33 samples from 11 individual batches of *E. coli* and 30 IBAT samples were analyzed by nuclear magnetic resonance (NMR) spectroscopy. The IBAT method reduces variance between different tubes of RM. The NMR spectra for these samples are nearly identical, with a very low variance (Fig. 2a). The variance here is due to extraction and quantification. In contrast, the variance between the 33 individual spectra is much larger, reflecting a combination of biological variance and technical variance. To quantify variance, we selected 19 metabolites that we could identify, were present in all the samples, were consistent between replicate measurements, with clear individual peaks enabling accurate quantification of individual metabolites. The coefficient of variation (CV – standard deviation/mean) was calculated separately for each metabolite within each group (Fig. 2b, Supp. Table 1). Similar to the overlaid NMR spectra (Fig. 2a), the CV was lower for IBAT generated samples (between 0.19 and 0.91) than for individual samples (0.36 to 1.26). Using the Fligner-Killeen^12^ test for homogeneity of variances for each of the selected metabolites showed significantly different variances between IBAT produced samples and individual batch samples (*p value* < 0.05) except for betaine (*p value* = 0.21).

**Figure 2:**
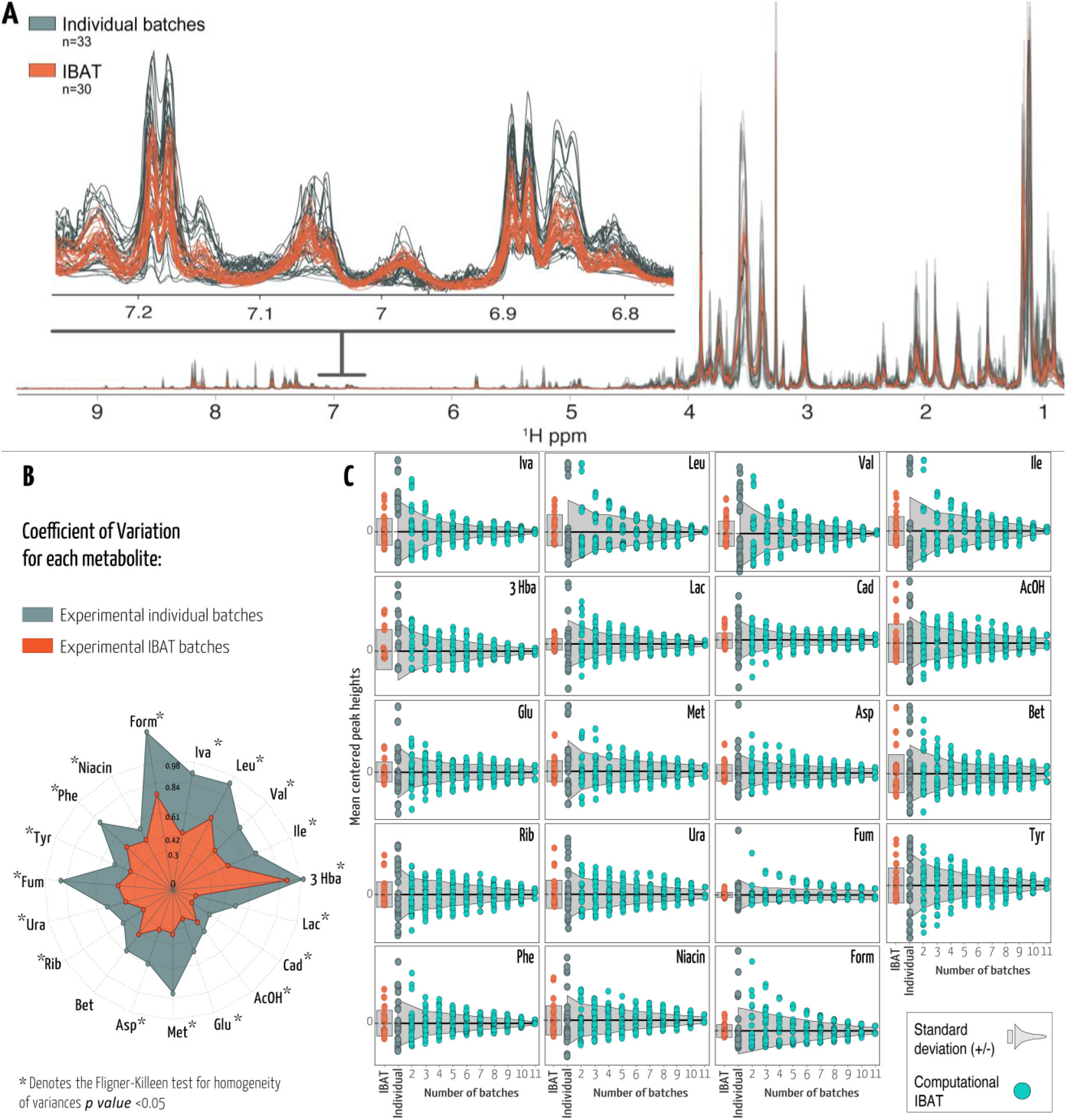
**A)** Untargeted full resolution ^1^H NMR profile of *E. coli* and spectral expansion between 6.8 and 7.2 ppm. NMR spectra in grey or orange correspond to IBAT or individual batches, respectively. **B)** Radial plot representing the coefficient of variation (CV) for annotated metabolites using the same colors. The length of spokes corresponds to the CV of each metabolite. **C)** Each data point represents the mean centered peak height in each sample. Experimental IBAT samples are depicted in orange and individual batches in grey. Cyan data points represent simulated metabolite peak heights per number of averaged batches. Light grey shaded areas represent +/− one standard deviation from the mean. Iva – isovalerate, Leu – leucine, Val – valine, Ile– isoleucine, 3 Hba – 3-hydroxybutyrate, Lac – lactate, Cad – cadaverine, AcOH – acetate, Glu– glutamate, Met – methionine, Asp – aspartate, Bet – betaine, Rib – ribose, Ura – uracil, Fum – fumarate, Tyr – tyrosine, Phe – phenylalanine, Niacin – nicotinic acid and Form – formate.

The IBAT process depends on pooling batches. We used the individual batch data to simulate the IBAT process. We generated 10 iterations for combining 2, 3, 4, 5, 6, 7, 8, 9, 10 and 11 individual batches to generate an IBAT compliant RM. We used the individual data to estimate the mean centered peak heights and respective standard deviations (Sd) for our 19 metabolites. The variance (10 iterations) decreases as the number of batches used increases (Fig. 2c). (Fig. 1 and Table.1). This is consistent with the predictions of Spearman-Brown.^13, 14^

### Production and application of a *C. elegans* PD1074 reference material

To create an IBAT *C. elegans* RM, we used a 2 L bioreactor and fed the worms the *E. coli* RM. Each batch of the bioreactor produced between 40-60 million mixed-stage worms. These were harvested and aliquoted into 20-30 tubes so that every tube contained 2 million worms. These were then frozen at −80 °C. After three bioreactor batches, we combined one aliquot from each batch for a total of 6 million batch-averaged worms. This was divided into 30 aliquots of *C. elegans* RM with 200 thousand worms each and refrozen until use (Fig. 3).

**Figure 3:**
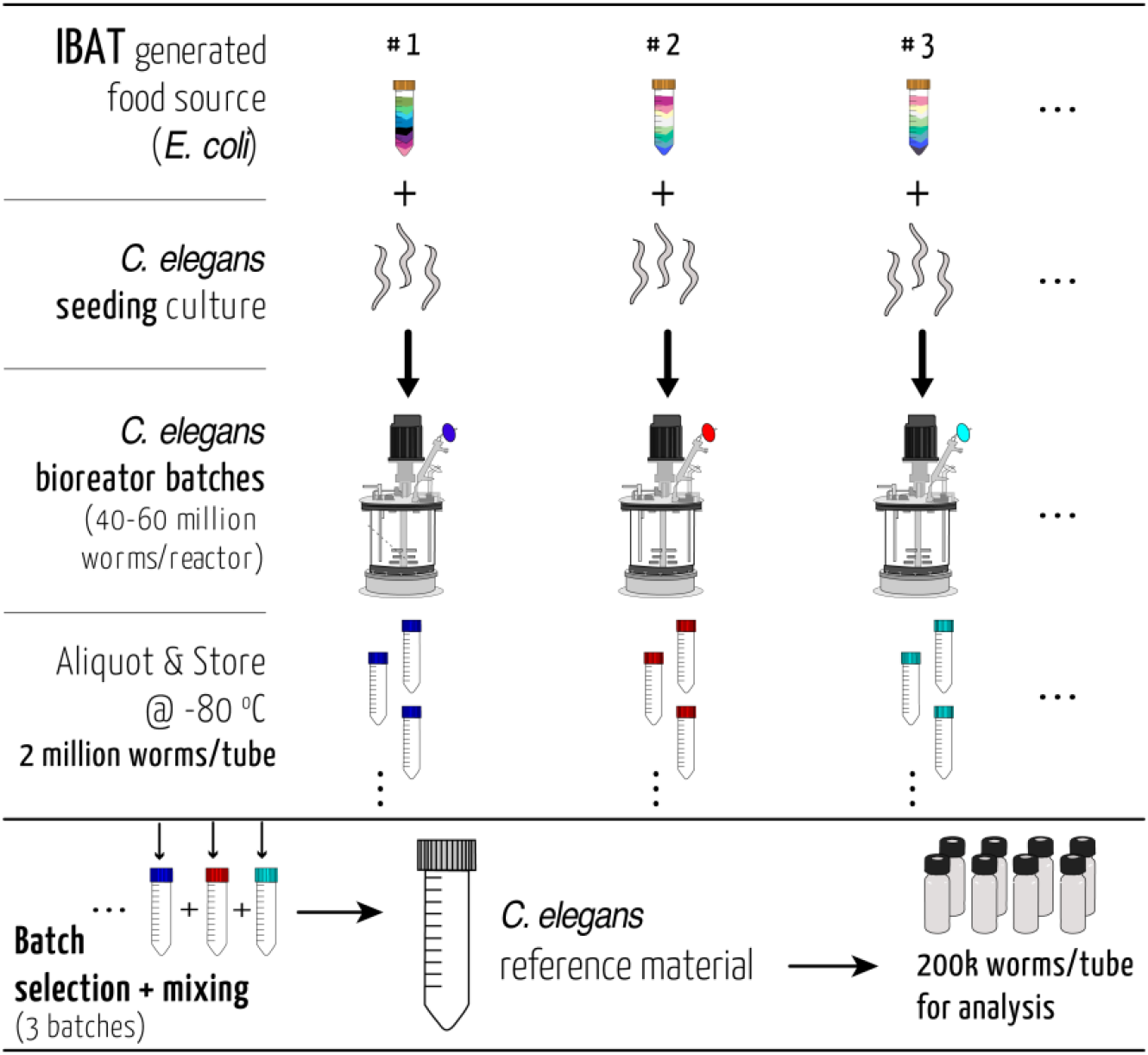
Schematic overview of the *C. elegans* reference material production. The reference strain PD1074 nematodes were seeded from cryo-preserved stocks and fed an *E. coli* RM (supplementary methods). Harvested material from each bioreactor was washed, aliquoted and stored. Aliquots from each reactor iteration were combined to produce a stable *C. elegans* reference material. This material can be divided into different sized aliquots according to the downstream application needs.

In a metabolomics experiment there are three main sources of variation: the sample material itself, the extraction, and data acquisition (supp. Fig 1). An experimental sample will encompass all three of those sources. The IBAT RM reduces the sample material variation, pooled samples average over both the sample variance and the extraction variation. We compared the *C. elegans* RM to 41 independent samples of the same strain (PD1074).^15^ These individual samples were prepared in three sets of two extraction blocks. For each set an equimolar pool was formed from all individual samples, for three pools. One *C. elegans* RM aliquot was included in each extraction block. In NMR data collection, one block was analyzed per each run. We selected 26 annotated features that were common to all samples and computed pairwise standardized Euclidean distances (SED) for each sample (Fig. 4). The distances between samples in the IBAT material reflect instrument variability (pools) and extraction variability. The distances between individual sample data include extraction and instrument variability but also sample variability. The mean and median distances, minimum and maximum values and sample distribution for each of these groups allow us to estimate the variability from these different sources of variation. The individual PD1074 samples, which include all three sources of variation, have the largest variability with mean values from 25.5 to 54.1 and the min/max of 8.19 and 66 (blocks 1 through 6 in Fig. 4). The IBAT samples, representing the extraction and technical variance, have a smaller range of mean distances (28.2 to 38.7) and min/max values of 21.3 and 44.8. As expected, the pooled individual PD1074 samples representing differences in the manual preparation and instrumentation between sets, have the smallest range with the respective boxplot bounds between 31.4 and 33.

**Figure 4:**
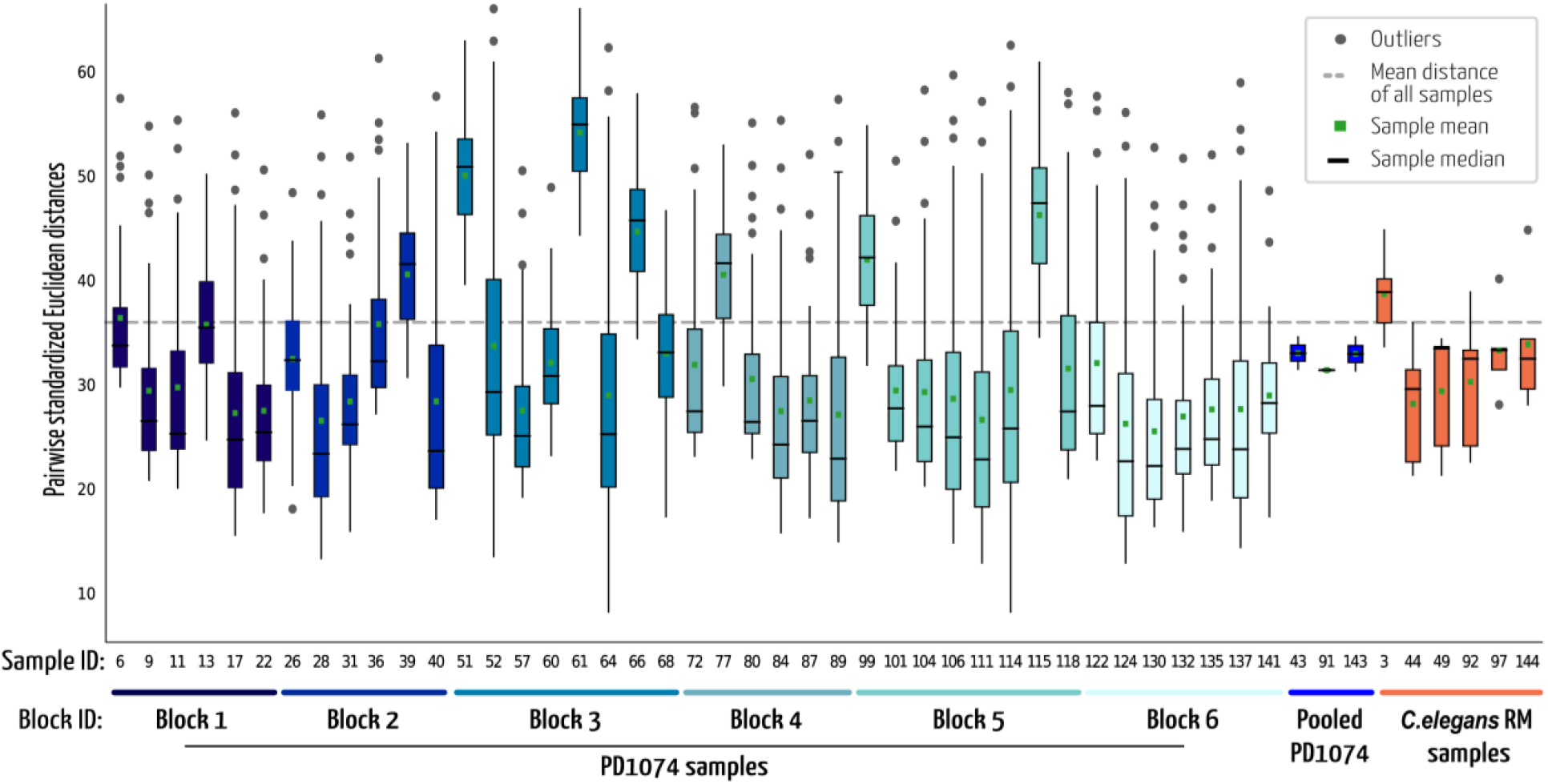
Boxplots of pair-wise standardized Euclidean distances. Each boxplot represents the distribution of distances from one sample to all the other samples of the same group. Mean and median distances for each sample are indicated by markers. Blue colored boxplots represent PD1074 samples that were processed in each block. The three pooled PD1074 samples were created from the samples in blocks 1+2, 3+4 and 5+6 respectively. *C. elegans* RM samples were generated using IBAT and processed alongside the PD1074 samples, one per each block.

## Discussion and Conclusion

The IBAT process reduces the growth and sampling contributions to variance by creating a common source of material from which homogeneous aliquots are produced. The advantage here is that instead of producing a single large batch, which will have its own challenges in achieving homogeneity, material is continuously generated over time, with each iteration using only small amounts of new material, thus capturing small changes over time while having minimal variance between experiments. This minimal variance can be theoretically predicted as a function of the number of distinct batches combined and the variance between the continuously produced material or, estimated from empirical data (Fig.2c) to take into account the overlap between iterations. The IBAT process is flexible and can be adjusted to production throughput, the type of material, the quantities produced the degree of variance reduction, and the metabolomics technology. We demonstrated this concept for two different types of matrices, *E. coli* and *C. elegans.* However, the method is general and can be applied to any biological matrix. In non-model systems studies it is common to use human plasma or urine or commercially available materials that are aliquoted from a single large batch and frozen. But when a batch runs out, shifting to a new external standard will often not be comparable to the prior standard. IBAT can be used by making pools from different batches of material as illustrated in Figure 1. New batches can be incorporated over time, and this will minimize the change in the RM over time. Similar strategies can be used with diverse applications such as plants or cultured mammalian cells for biotherapeutics. In these scenarios the main issue is minimizing the freeze thaw cycles and so, the size of the initial aliquots for future blending must be planned.

A RM of the same biological matrix as the study samples together with a carefully planned experimental design can be used to determine the magnitude and variance in the extraction, a major source of variation in metabolomics experiments.^3, 16^ It can also facilitate comparison among separate experiments. The IBAT can then be used to separate the extraction variance from the sample-to-sample variance in the individually grown and processed samples, as demonstrated here. The individual *C. elegans* samples are genetically identical to the RM. Variance in metabolite intensities were larger as a result of sample variation during growth, handling, storage and sampling, added to the technical variation in extraction and data acquisition. The pooled individual PD1074 sample minimizes the sample and extraction variance by averaging over both samples and extractions and reflects only the variation in the analytical measurement (which is low for NMR) and the pooling strategy. By processing the experimental replicates and RM aliquots, one can independently estimate the contribution of the metabolite extraction step to individual metabolite variation. The IBAT *C. elegans* RM samples can be used to estimate variance due to extraction. We find that 40% of the total variance as estimated by the variation between individually grown, extracted and quantified samples is due to extraction variance and analysis and of that variance ~15% is due to technical variation.

IBAT increases the efficacy of QA/QC and is expected to improve the performance of biological reference materials by allowing estimation of process-derived variance including facilitating studies across multiple labs. Finally, the cost of using an IBAT process should be lower than acquiring a single large batch of reference material thus enabling labs to amortize the process over time while maintaining the stability of the material and facilitating comparison of experiments conducted months or years apart.

## Methods

### *E. coli* individual batches production and storage

In order to produce a stable and consistent *C. elegans* food source, batches of *E. coli* HT115 were grown in bioreactors (Biostat, Sartorius) using standardized protocols (Supplementary methods). A total of 11 batches were produced and each batch was divided into approximately 60-90 aliquots, flash frozen and stored at −80 °C. Each aliquot is comprised of 2 mL of bacterial suspension (1 g of wet bacterial paste and OD_600_ ranging from 17.5 to 24).

### NMR sample preparation of *E. coli* IBAT and individual batches

All 33 individual batch samples and 30 IBAT generated samples were prepared for NMR analysis. Approximately 200 μL of 0.7 mm silica beads (BioSpec products) were added to each of the 63 samples. These were homogenized at 1800 rpm for 300s (FastPrep 96 - MPBIO) and centrifuged at 20,000 × G for 15 minutes. From each sample, 450 μL of supernatant were transferred to a new tube and 150 μL of deuterated water were added (D_2_O, D, 99.9%, Cambridge Isotope Laboratories). Each sample was vortex-mixed for 1 min before transferring into 5 mm SampleJet NMR tubes. Details of NMR acquisition and spectra processing can be found in the supplementary methods.

### NMR sample preparation of *C. elegans* samples

For the NMR analysis six IBAT RM aliquots were prepared for alongside 41 individual samples of the *C. elegans* strain PD1074 that were grown according to our previously published method.^15^ Each of these samples contained approximately 200,000 nematodes. All samples were previously flash frozen and then lyophilized until dry. Approximately 200 μL of 1 mm Zirconia beads (BioSpec products) were added to each dry sample and homogenized at 1800 rpm for a total of 270 seconds (FastPrep 96 - MPBIO). The samples were then delipidated by adding 1 mL of cold (−20 °C) isopropanol (Optima, LC/MS Grade, Fisher Scientific) and left overnight (12 hours) at −20 °C after a 20 min resting period at room temperature. The supernatant was removed after being centrifuged for 30 min at 20,000 × G and 1 mL of cold (4 °C) 80/20 methanol/water (Optima, LC/MS Grade, Fisher Scientific) was added to the remaining contents. The tubes were shaken for 30 min at 4 °C and centrifuged at 20,000 × G for 30 minutes. The methanol/water supernatant was transferred to new tubes and these were vacuum dried using a CentriVac benchtop vacuum concentrator (Labconco). The extracts were reconstituted in 45 μl of deuterated (D_2_O, D, 99.9%, Cambridge Isotope Laboratories) 100 mM sodium phosphate buffer (mono- and dibasic; Fisher BioReagents) containing 0.11 mM of the internal standard DSS (sodium 2,2-dimethyl-2-silapentane-5-sulfonate, D6, 98%; Cambridge Isotope Laboratories) at pH 7.0 and vortex mixed for <1 min prior to transfer into 1.7 mm SampleJet NMR tubes. The three pooled PD1074 samples were created by adding together 6 μl from the samples in each NMR run (12, 14 and 15 samples respectively), after having been reconstituted in the internal standard containing NMR solvent. Details of NMR acquisition and spectra processing can be found in the supplementary methods.

### Data analysis

Following acquisition and processing, spectra were imported into Matlab programing software (MATLAB, MathWorks, R2019a). Using a metabolomics toolbox developed in-house and freely available (https://github.com/artedison/Edison_Lab_Shared_Metabolomics_UGA) the following was carried out: plotting, referencing, baseline correction, alignment (CCOW^17^) and solvent peaks removal. Feature detection (peak picking) was automated using a combination of an in-house peak picking function and binning algorithm^18^ to extract peak heights. Data were exported for Bland-Altman analysis, to select features that are in agreement between its replicates (cut-offs used: sample flag of 0.2, feature flag of 0.05 and residual of 3), and pairwise Standardized Euclidean Distances (SED) analysis using the SouthEast Center for Integrated Metabolomics Tools (SECIMTools)^19^. Coefficient of variation calculations (CV), variance, %variance and Fligner-Killeen test were carried out in Matlab.

### Data availability

All raw and processed data, along with detailed experimental NMR and data analysis methods, will be available upon processing at Metabolomics Workbench (https://www.metabolomicsworkbench.org).

## Acknowledgement

Research reported in this publication was supported by the National Institutes of Health under Award Number 1U2CES030167-01. The authors would like to thank Dr. David Blum and Ron Garrison from the Bio-expression and Fermentation Facility at the University of Georgia for training and advice using the Bioreactors and Pamela Kirby at the Edison Lab for assistance with material handling and storage logistics.

## References

1. Cochran, W. G. a. C., G.M., Experimental Design. 2nd edition ed.; New York, 1957.

2. Dunn, W. B.; Broadhurst, D. I.; Edison, A.; Guillou, C.; Viant, M. R.; Bearden, D. W.; Beger, R. D., Quality assurance and quality control processes: summary of a metabolomics community questionnaire. Metabolomics 2017, 13 (5).

3. Broadhurst, D.; Goodacre, R.; Reinke, S. N.; Kuligowski, J.; Wilson, I. D.; Lewis, M. R.; Dunn, W. B., Guidelines and considerations for the use of system suitability and quality control samples in mass spectrometry assays applied in untargeted clinical metabolomic studies. Metabolomics 2018, 14 (6), 72.

4. Paulovich, A. G.; Billheimer, D.; Ham, A. J.; Vega-Montoto, L.; Rudnick, P. A.; Tabb, D. L.; Wang, P.; Blackman, R. K.; Bunk, D. M.; Cardasis, H. L.; Clauser, K. R.; Kinsinger, C. R.; Schilling, B.; Tegeler, T. J.; Variyath, A. M.; Wang, M.; Whiteaker, J. R.; Zimmerman, L. J.; Fenyo, D.; Carr, S. A.; Fisher, S. J.; Gibson, B. W.; Mesri, M.; Neubert, T. A.; Regnier, F. E.; Rodriguez, H.; Spiegelman, C.; Stein, S. E.; Tempst, P.; Liebler, D. C., Interlaboratory study characterizing a yeast performance standard for benchmarking LC-MS platform performance. Mol Cell Proteomics 2010, 9 (2), 242–54.

5. Phinney, K. W.; Ballihaut, G.; Bedner, M.; Benford, B. S.; Camara, J. E.; Christopher, S. J.; Davis, W. C.; Dodder, N. G.; Eppe, G.; Lang, B. E.; Long, S. E.; Lowenthal, M. S.; McGaw, E. A.; Murphy, K. E.; Nelson, B. C.; Prendergast, J. L.; Reiner, J. L.; Rimmer, C. A.; Sander, L. C.; Schantz, M. M.; Sharpless, K. E.; Sniegoski, L. T.; Tai, S. S.; Thomas, J. B.; Vetter, T. W.; Welch, M. J.; Wise, S. A.; Wood, L. J.; Guthrie, W. F.; Hagwood, C. R.; Leigh, S. D.; Yen, J. H.; Zhang, N. F.; Chaudhary-Webb, M.; Chen, H.; Fazili, Z.; LaVoie, D. J.; McCoy, L. F.; Momin, S. S.; Paladugula, N.; Pendergrast, E. C.; Pfeiffer, C. M.; Powers, C. D.; Rabinowitz, D.; Rybak, M. E.; Schleicher, R. L.; Toombs, B. M.; Xu, M.; Zhang, M.; Castle, L., Development of a Standard Reference Material for metabolomics research. Anal Chem 2013, 85 (24), 11732–8.

6. Simon-Manso, Y.; Lowenthal, M. S.; Kilpatrick, L. E.; Sampson, M. L.; Telu, K. H.; Rudnick, P. A.; Mallard, W. G.; Bearden, D. W.; Schock, T. B.; Tchekhovskoi, D. V.; Blonder, N.; Yan, X.; Liang, Y.; Zheng, Y.; Wallace, W. E.; Neta, P.; Phinney, K. W.; Remaley, A. T.; Stein, S. E., Metabolite profiling of a NIST Standard Reference Material for human plasma (SRM 1950): GC-MS, LC-MS, NMR, and clinical laboratory analyses, libraries, and web-based resources. Anal Chem 2013, 85 (24), 11725–31.

7. Han, W.; Li, L., Evaluating and minimizing batch effects in metabolomics. Mass Spectrom Rev 2020.

8. Peng, J.; Chen, Y. T.; Chen, C. L.; Li, L., Development of a universal metabolome-standard method for long-term LC-MS metabolome profiling and its application for bladder cancer urine-metabolite-biomarker discovery. Anal Chem 2014, 86 (13), 6540–7.

9. Bunk, D. M., Design considerations for proteomic reference materials. Proteomics 2010, 10 (23), 4220–5.

10. Metrology, J. C. f. G. i., International vocabulary of metrology - Basic and general concepts and associated terms. VIM 2012, (200).

11. Beger, R. D.; Dunn, W. B.; Bandukwala, A.; Bethan, B.; Broadhurst, D.; Clish, C. B.; Dasari, S.; Derr, L.; Evans, A.; Fischer, S.; Flynn, T.; Hartung, T.; Herrington, D.; Higashi, R.; Hsu, P. C.; Jones, C.; Kachman, M.; Karuso, H.; Kruppa, G.; Lippa, K.; Maruvada, P.; Mosley, J.; Ntai, I.; O’Donovan, C.; Playdon, M.; Raftery, D.; Shaughnessy, D.; Souza, A.; Spaeder, T.; Spalholz, B.; Tayyari, F.; Ubhi, B.; Verma, M.; Walk, T.; Wilson, I.; Witkin, K.; Bearden, D. W.; Zanetti, K. A., Towards quality assurance and quality control in untargeted metabolomics studies. Metabolomics 2019, 15 (1), 4.

12. Conover, W. J.; Johnson, M. E.; Johnson, M. M., A Comparative Study of Tests for Homogeneity of Variances, with Applications to the Outer Continental Shelf Bidding Data. Technometrics 1981, 23 (4), 351–361.

13. Brown, W., Some Experimental Results in the Correlation of Mental Abilities1. British Journal of Psychology, 1904-1920 1910, 3 (3), 296–322.

14. Spearman, C., Correlation Calculated from Faulty Data. British Journal of Psychology, 1904-1920 1910, 3 (3), 271–295.

15. Amanda O. Shaver, G. J. G., Pamela S. Kirby, Erik Andersen, Arthur S. Edison, Culture and assay of Large-Scale Mixed Stage Caenorhabditis elegans Population. JOVE - J. Vis. Exp 2020, e61453.

16. Liu, Q.; Walker, D.; Uppal, K.; Liu, Z.; Ma, C.; Tran, V.; Li, S.; Jones, D. P.; Yu, T., Addressing the batch effect issue for LC/MS metabolomics data in data preprocessing. Sci Rep 2020, 10 (1), 13856.

17. Tomasi, G.; van den Berg, F.; Andersson, C., Correlation optimized warping and dynamic time warping as preprocessing methods for chromatographic data. Journal of Chemometrics 2004, 18 (5), 231–241.

18. Sousa, S. A. A.; Magalhães, A.; Ferreira, M. M. C., Optimized bucketing for NMR spectra: Three case studies. Chemometrics and Intelligent Laboratory Systems 2013, 122, 93–102.

19. Kirpich, A. S.; Ibarra, M.; Moskalenko, O.; Fear, J. M.; Gerken, J.; Mi, X.; Ashrafi, A.; Morse, A. M.; McIntyre, L. M., SECIMTools: a suite of metabolomics data analysis tools. BMC Bioinformatics 2018, 19 (1), 151.

