## Supplementary material for "Long-Term Metabolomics Reference Material": supp. methods

**Supplementary methods:**

Bioreactor production of *Escherichia coli*:

Cultures of *E. coli* were started from frozen stocks kept at -80 $^oC$. A streak plate with LB media and agar (LB broth, Miller – Novagen, agar, Bacto BD) was made under standard aseptic conditions and incubated for 32 hours at 37 $^oC$. A single colony was transferred to 20 mL of Terrific Broth (TB - Fisher BioReagents) and placed in a shaker incubator for 24 hours at 37 $^oC$ and 250 rpm. A contamination control LB plate was then streaked under aseptic conditions and incubated overnight. This liquid culture was the starting inoculum for a bioreactor (Biostat A, Sartorius) containing 2 L of TB with 30 mL of glycerol (EMD millipore). The bioreactor was under automated control of temperature, dissolved oxygen, pH (37 $^oC$, 30% and 7.5 respectively) and constant mixing (500 rpm). After 42 hours growth an $OD_{600}$ measurement was taken, and the bioreactor harvested into 500 mL centrifuge bottles and centrifuged for 30 min at 10,000 x G. The supernatant was discarded and the pellet process repeated two more times with deionized water and finally the centrifuged pellets were combined, weighed and reconstituted in M9 minimal media[1] to a concentration of 0.5 g/mL. Aliquots from this material were then made so that each tube contains 1 g (wet weight) of material by an automated pipetting robot (Andrew Robot – Andrew Alliance), flash frozen in liquid Nitrogen and stored at -80 $^oC$.

Making a stable food source for *C. elegans* growth:

Individual batches of *E. coli* were produced as described above. Six aliquots from 10 different individual bacterial batches were thawed on ice and pooled together as substrate for one *C. elegans* batch. For optimal *C. elegans* growth, a total of 60 bacterial aliquots were required to achieve a ratio of 3% (w/v)[2] of substrate to volume of media in a 2 L bioreactor (Biostat, Sartorius). IBAT was used to create two additional batches of food, each containing 60 aliquots from 10 batches where, for each iteration, aliquots from one individual batch were removed, and new individual batch aliquots added.

Growing *C. elegans* in bioreactors:

Similar to the *E. coli* bioreactor process a starting inoculum of *C. elegans* was first made. This was a population of worms collected from a large scale culture plate as described previously.[3]
Approximately 2 million worms were washed with M9 media and added to the bioreactor (Biostat A, Sartorius) containing 2 L of K-media[1] and the stable food source created above. The Bioreactor was under automated control of temperature, dissolved oxygen, pH (20 $^oC$, 10% and 7 respectively) and constant mixing (150 rpm). Two daily $OD_{600}$ measurements were taken to monitor the amount of available food and the nematodes counted under the microscope to account for overcrowding. The bioreactor was harvested when food was below 0.5% w/v (calculated from $OD_{600}$ measurements) and/or nematode density was above 30,000 individuals/mL. The harvested culture was then divided into 500 mL centrifuge bottles and centrifuged for 20 min at 5,000 x G

and 4 °C. The supernatant discarded, and the wash process repeated two more times with M9 media and a final reconstitution with deionized water. The contents of each bottle were combined, and three 1 mL aliquots taken to count the number of nematodes[3]. The material was then aliquoted into 15 mL centrifuge tubes by an automated pipetting robot (Andrew Robot – Andrew Alliance) each containing approximately 2,000,000 nematodes, flash frozen in liquid Nitrogen and stored at -80 °C.

NMR data acquisition and processing:

One-dimensional $^{1}$H NMR spectra were acquired using moesypr1d with pre-saturation during relaxation delay and mixing time on an Avance III HD 600 MHz Bruker NMR spectrometer equipped with a TCI cryoprobe and a Bruker SampleJet autosampler cooled to 5.6 °C. During acquisition, 32,768 complex datapoints were collected for the FID, using 64 scans with 4 additional dummy scans. The spectral width was 20 ppm. A Fourier transform (FT), a polynomial baseline correction of order 2, a 0.5 Hz line broadening and phase correction were applied to each spectrum using NMRPipe processing software.[4]

Two-dimensional $^{1}$H-$^{1}$H total correlation spectroscopy (TOCSY- dipsi2esfbgpph), $^{1}$H-$^{13}$C heteronuclear single quantum correlation (HSQC - hsqcedetgpsisp2.3) and $^{1}$H-$^{13}$C HSQC–total correlation spectroscopy (HSQC–TOCSY - hsqcdietgpsisp.2) experiments were collected on both *C. elegans* and *E. coli* samples for metabolite identification. During acquisition, all three experiments were collected for 32 scans and an additional 16 dummy scans, with 512 and 1,024 datapoints recorded on the direct and indirect dimensions respectively and, a spectral width of 200 ppm for $^{13}$C and 12 ppm for $^{1}$H. A 90ms mixing time was used for both HSQC-TOCSY and TOCSY experiments. All spectral processing was carried out using NMRPipe [5].

Compound identification/database matching:

All two-dimensional experiments were used for spectral matching against the BBiorefcode library using COLMARm[6] and a chemical shift cutoff of 0.03 and 0.3 ppm for $^{1}$H and $^{13}$C respectively. Metabolites that could be quantified without overlap and were consistent between replicates in their respective 1D $^{1}$HNMR spectra were selected to be identified. From the *E. coli* samples 19 features were annotated to metabolites and 26 in the *C. elegans* samples. A confidence level ranging from 1 to 5 (Supplementary Table 1), was assigned to each metabolite as described elsewhere [7]. Briefly this scale is defined as: (1) putatively characterized compound, (2) matched to reported 1D spectra, (3) matched to reported HSQC spectra, (4) matched to reported HSQC and HSQC-TOCSY spectra, and (5) validated by spiking putative compound into sample.

### Supplementary figures:

Supplementary Table 1a: Table of metabolites isolated features that were common to all for *E. coli* samples

| | | | Individual batches | | | IBAT batches | | |
|---|---|---|---|---|---|---|---|---|
| Compound name (COLMARm) | Identification level | ppm_1D | Mean | Standard deviation | CV (mean/std) | Mean | Standard deviation | CV (mean/std) |
| 'Isovaleric_acid_1 | 4 | 0.8903 | 6.68E+07 | 6.20E+07 | 0.9281 | 5.83E+07 | 2.64E+07 | 0.4528 |
| 'Leucine_1 | 4 | 0.9375 | 8.25E+07 | 7.83E+07 | 0.9501 | 5.27E+07 | 3.42E+07 | 0.6494 |
| 'L_Valine_1 | 4 | 0.9695 | 5.69E+07 | 4.14E+07 | 0.7278 | 3.37E+07 | 1.56E+07 | 0.4622 |
| 'L_Isoleucine_1 | 4 | 0.9929 | 3.58E+07 | 2.57E+07 | 0.7185 | 2.25E+07 | 1.09E+07 | 0.4865 |
| '3_Hydroxybutyrate_1 | 3 | 1.1741 | 3.91E+07 | 4.03E+07 | 1.0322 | 2.34E+07 | 2.14E+07 | 0.9116 |
| 'Lactic_acid_1 | 4 | 1.2943 | 3.10E+07 | 1.60E+07 | 0.5143 | 2.04E+07 | 4.04E+06 | 0.1978 |
| 'Cadaverine_1 | 3 | 1.4481 | 9.72E+07 | 3.49E+07 | 0.3594 | 7.48E+07 | 1.45E+07 | 0.1943 |
| 'Acetic_acid_1 | 3 | 1.8941 | 1.04E+08 | 4.31E+07 | 0.4136 | 8.88E+07 | 2.86E+07 | 0.3220 |
| 'L_Glutamic_acid_1 | 3 | 2.3281 | 4.45E+07 | 2.31E+07 | 0.5194 | 3.61E+07 | 8.71E+06 | 0.2413 |
| 'L_Methionine_1 | 4 | 2.6190 | 8.95E+06 | 7.27E+06 | 0.8119 | 7.33E+06 | 2.58E+06 | 0.3524 |
| 'D_Aspartate_1 | 3 | 2.6564 | 7.67E+06 | 4.70E+06 | 0.6119 | 6.16E+06 | 2.07E+06 | 0.3360 |
| 'Betaine_1 | 4 | 3.2377 | 8.80E+08 | 5.33E+08 | 0.6061 | 8.56E+08 | 3.54E+08 | 0.4138 |
| 'D_Ribose_2 | 4 | 4.8944 | 8.61E+06 | 4.07E+06 | 0.4726 | 6.62E+06 | 1.87E+06 | 0.2830 |
| 'Uracil_1 | 4 | 5.7637 | 8.69E+06 | 4.68E+06 | 0.5389 | 6.75E+06 | 2.69E+06 | 0.3981 |
| 'Fumaric_acid_1 | 3 | 6.4871 | 4.88E+05 | 4.34E+05 | 0.8898 | 1.69E+05 | 7.51E+04 | 0.4453 |
| 'L_Tyrosine_1 | 3 | 7.1577 | 6.45E+06 | 3.25E+06 | 0.5043 | 4.67E+06 | 1.74E+06 | 0.3716 |
| L_Phenylalanine_1 | 4 | 7.4040 | 8.04E+06 | 6.34E+06 | 0.7890 | 7.04E+06 | 3.56E+06 | 0.5057 |
| 'Nicotinic_acid_1 | 4 | 8.2322 | 1.24E+06 | 6.80E+05 | 0.5498 | 8.79E+05 | 4.00E+05 | 0.4553 |
| 'Formate_1 | 3 | 8.4308 | 3.37E+06 | 4.24E+06 | 1.2577 | 1.50E+06 | 1.15E+06 | 0.7647 |

Supplementary Table 1b: Table of metabolites isolated features that were common to all for *C. elegans* samples.

| Compound name (COLMARm) | Identification level | ppm_1D | PD1074 Mean | PD1074 SD | IBAT Mean | IBAT SD | PoolQC Mean | PoolQC SD |
|---|---|---|---|---|---|---|---|---|
| 'AMP_sulfate_1' | 3 | 8.2697 | 1.05E+10 | 6.43E+09 | 6.52E+09 | 6.18E+08 | 1.18E+10 | 2.73E+09 |
| 'Benzoate_1' | 3 | 7.4853 | 2.41E+09 | 1.41E+09 | 3.11E+09 | 1.59E+09 | 2.50E+09 | 9.87E+08 |
| 'L_Phenylalanine_1' | 4 | 7.4387 | 2.31E+09 | 1.73E+09 | 4.06E+09 | 1.70E+09 | 2.18E+09 | 2.89E+08 |
| 'L_Tyrosine_1' | 4 | 6.8948 | 3.04E+09 | 2.04E+09 | 6.30E+09 | 1.34E+09 | 3.34E+09 | 5.80E+08 |
| 'UDP_GlcNAc_1' | 3 | 5.5317 | 1.63E+09 | 8.80E+08 | 1.20E+09 | 1.47E+08 | 1.60E+09 | 1.93E+08 |
| 'Allantoin_1' | 3 | 5.3865 | 4.11E+09 | 2.86E+09 | 3.54E+09 | 6.51E+08 | 4.40E+09 | 9.41E+08 |
| 'D_Trehalose_1' | 4 | 5.188 | 6.04E+10 | 3.54E+10 | 1.47E+11 | 1.23E+10 | 6.25E+10 | 1.80E+09 |
| 'D_Glucose_1' | 4 | 4.6492 | 5.13E+09 | 2.73E+09 | 8.45E+09 | 1.31E+09 | 5.28E+09 | 8.57E+08 |
| 'Betaine_1' | 4 | 3.2672 | 3.15E+11 | 2.75E+11 | 1.08E+12 | 1.29E+11 | 3.82E+11 | 1.63E+11 |
| 'Lysine_1' | 4 | 3.054 | 2.67E+10 | 1.53E+10 | 1.50E+10 | 2.18E+09 | 2.99E+10 | 3.78E+09 |
| 'L_Asparagine_1' | 3 | 2.9527 | 4.40E+09 | 2.71E+09 | 1.62E+09 | 1.27E+09 | 5.37E+09 | 4.41E+08 |
| 'D_Aspartate_1' | 3 | 2.7968 | 7.35E+09 | 4.32E+09 | 3.63E+09 | 1.71E+09 | 7.31E+09 | 1.16E+09 |
| 'Allocystathionine_1' | 3 | 2.741 | 1.47E+10 | 8.99E+09 | 2.32E+09 | 2.54E+08 | 1.48E+10 | 1.07E+09 |
| 'Succinic_acid_1' | 3 | 2.4119 | 2.78E+11 | 2.13E+11 | 2.44E+11 | 2.86E+10 | 3.13E+11 | 6.05E+10 |
| 'L_Glutamic_acid_1' | 3 | 2.3585 | 5.98E+10 | 4.13E+10 | 7.65E+10 | 1.06E+10 | 6.42E+10 | 1.16E+10 |
| '2_Aminoadipic_acid_1' | 3 | 2.2476 | 1.36E+10 | 1.17E+10 | 1.21E+10 | 1.87E+09 | 1.18E+10 | 2.40E+08 |
| 'N_acetyl_putrescine_1' | 4 | 1.9906 | 7.40E+10 | 7.02E+10 | 8.24E+10 | 9.18E+09 | 7.60E+10 | 1.19E+10 |
| 'Acetic_acid_1' | 3 | 1.9208 | 6.15E+10 | 4.26E+10 | 1.59E+11 | 2.29E+10 | 6.57E+10 | 1.61E+10 |
| 'Putrescine_1' | 4 | 1.7771 | 2.67E+10 | 1.51E+10 | 1.08E+10 | 4.98E+09 | 2.67E+10 | 2.57E+09 |
| 'Alanine_1' | 4 | 1.4761 | 3.46E+11 | 2.29E+11 | 5.59E+11 | 1.09E+11 | 3.65E+11 | 5.96E+10 |
| 'Lactic_acid_1' | 4 | 1.3238 | 1.02E+11 | 8.79E+10 | 1.81E+11 | 2.13E+10 | 1.11E+11 | 4.34E+09 |
| 'Propionic_acid_1' | 3 | 1.0672 | 6.16E+09 | 4.94E+09 | 3.63E+10 | 4.89E+09 | 9.85E+09 | 5.60E+09 |
| 'L_Isoleucine_1' | 4 | 1.0192 | 1.07E+10 | 5.97E+09 | 3.51E+10 | 1.82E+10 | 1.30E+10 | 2.67E+09 |
| 'L_Valine_1' | 4 | 0.98857 | 1.82E+10 | 9.89E+09 | 6.06E+10 | 2.87E+10 | 2.01E+10 | 1.80E+09 |
| 'Leucine_1' | 4 | 0.97397 | 1.95E+10 | 1.10E+10 | 5.78E+10 | 2.60E+10 | 2.17E+10 | 1.86E+09 |
| 'L_Isoleucine_1' | 4 | 0.94007 | 3.37E+10 | 1.35E+10 | 6.01E+10 | 2.13E+10 | 3.55E+10 | 3.36E+09 |
| 'Pantothenate_1' | 3 | 0.89793 | 2.42E+10 | 1.28E+10 | 4.53E+10 | 8.29E+09 | 2.86E+10 | 3.00E+09 |

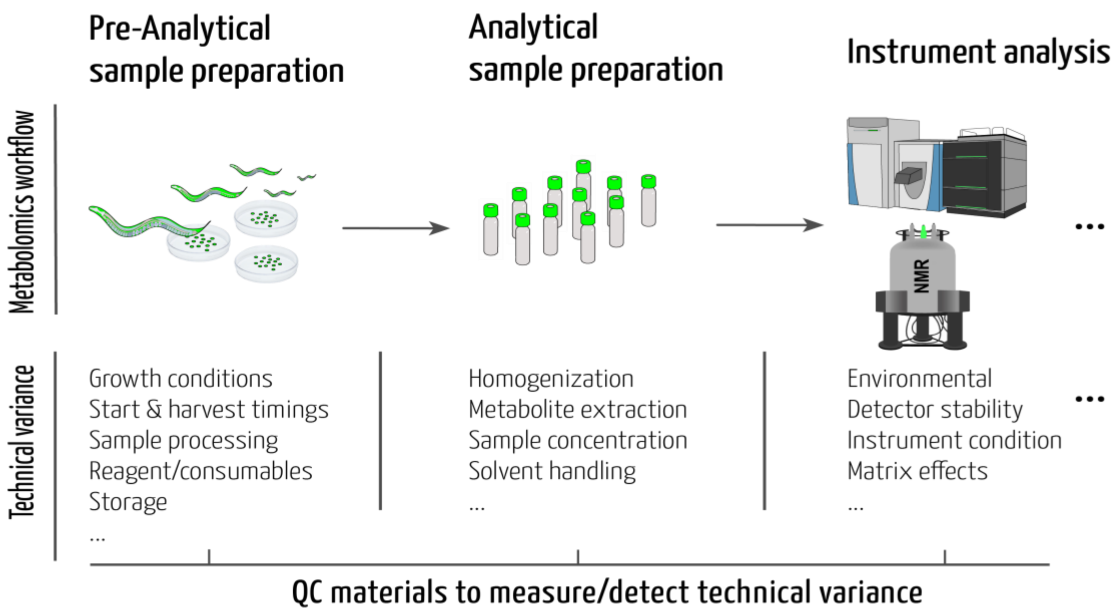


**Supplementary Figure 1:** General metabolomics workflow from sample generation to instrument analysis. Non-exhaustive examples of pre-analytical technical variance at each step of the metabolomics process.

### References:


1. Steiernagle, T. Maintenance of C. elegans - Wormbook http://www.wormbook.org/.
2. Kaplan, F.; Srinivasan, J.; Mahanti, P.; Ajredini, R.; Durak, O.; Nimalendran, R.; Sternberg, P. W.; Teal, P. E.; Schroeder, F. C.; Edison, A. S.; Alborn, H. T., Ascaroside expression in Caenorhabditis elegans is strongly dependent on diet and developmental stage. *PLoS One* **2011,** *6* (3), e17804.
3. Amanda O. Shaver, G. J. G., Pamela S. Kirby, Erik Andersen, Arthur S. Edison, Culture and assay of Large-Scale Mixed Stage Caenorhabditis elegans Population. *JOVE - J. Vis. Exp* **2020,** *e61453*.
4. Delaglio, F.; Grzesiek, S.; Vuister, G. W.; Zhu, G.; Pfeifer, J.; Bax, A., NMRPipe: a multidimensional spectral processing system based on UNIX pipes. *J Biomol NMR* **1995,** *6* (3), 277-93.
5. Delaglio, F.; Grzesiek, S.; Vuister, G. W.; Zhu, G.; Pfeifer, J.; Bax, A., NMRPipe: A multidimensional spectral processing system based on UNIX pipes. *J. Biomol. NMR* **1995,** *6* (3), 277-293.
6. Bingol, K.; Li, D. W.; Zhang, B.; Bruschweiler, R., Comprehensive metabolite identification strategy using multiple two-dimensional NMR spectra of a complex mixture implemented in the COLMARm web server. *Anal. Chem.* **2016,** *88* (24), 12411-12418.
7. Walejko, J. M.; Chelliah, A.; Keller-Wood, M.; Gregg, A.; Edison, A. S., Global metabolomics of the placenta reveals distinct metabolic profiles between maternal and fetal placental tissues following delivery in non-labored women. *Metabolites* **2018,** *8* (1).
